# Vitrification-warming delays preimplantation development and impairs mitochondrial function and cytoplasmic lattices integrity in mouse embryos

**DOI:** 10.1101/2025.05.05.652170

**Authors:** Mariana T. Barroso, Jose A. Rodríguez Muñoz, Viola Sjöström, Konstantina Dindini, Jian Zhao, Kenny A. Rodriguez-Wallberg, Arturo Reyes-Palomares

## Abstract

Vitrification and warming of human embryos have become standardized procedures in assisted reproduction over the past two decades. Although generally considered safe, their full impact on embryo development remains unclear. Epidemiological studies have raised concerns about differences in birth weight and long-term health outcomes between newborns resulting from fresh versus frozen embryo transfers. In this study, we used mouse embryos to investigate the impact of vitrification and warming on developmental kinetics, mitochondrial function, and cytoplasmic lattice integrity. Time-lapse imaging revealed significant developmental delays across all preimplantation stages in vitrified embryos. Additionally, mitochondrial distribution, volume, and membrane potential exhibited signs of impairment. Ultrastructural analysis identified damage such as ruptured mitochondrial membranes, disrupted cytoplasmic lattices during early cell divisions, and underdeveloped mitochondrial cristae at the blastocyst stage. We hypothesize embryo vitrification and warming disrupt mitochondrial function and destabilize cytoplasmic lattice integrity, ultimately contributing to developmental delays in preimplantation embryos.

## INTRODUCTION

Cryopreservation has revolutionized assisted reproduction, particularly with the implementation of the “freeze-all” policy, which involves storing embryos for future use instead of immediate transfer. This strategy helps mitigate risks of ovarian hyperstimulation and improves success rates in cases involving endometrial anomalies or requiring preimplantation genetic testing^1,2^. Additionally, embryo cryopreservation is increasingly used for social freezing, allowing women to preserve their fertility for later childbearing^3^. According to the European Society of Human Reproduction and Embryology (ESHRE), by 2014, frozen embryo transfers (FETs) accounted for over 50% of all embryo transfers in some European countries^4^.

Despite its widespread adoption, the rapid expansion of cryopreservation procedures has outpaced our ability to fully understand their impact on embryo development and offspring health. Recent population-based registry studies have raised concerns about potential risks associated with embryo cryopreservation, including (1) an increased risk of infant mortality (from birth up to 1 year of life) in singletons resulting from FETs^5^, (2) an increased risk of obstetrical complications in singletons births following FET of two embryos^6^, (3) a higher incidence of childhood cancer^7,8^, and (4) imprinting disorders in children born from FETs^9^. A notable strength of these studies is their focus on singleton births, minimizing confounders from multiple pregnancies and allowing clearer comparisons between ART and natural conception.

Over the past decade, vitrification has replaced slow freezing as the preferred method for embryo cryopreservation, offering improved success rates for frozen versus fresh embryo transfers^10–12^. First applied successfully to human embryos in the early 1980s, slow freezing often led to intracellular ice crystal formation, causing cellular damage and reduced survival rates^13^. Vitrification, introduced in the lates 1990s, uses ultra-rapid cooling to avoid ice formation, dramatically improving post-thaw viability^14,15^. Techniques such as the Cryotop method, developed by Kuwayama, further refined vitrification by minimizing solution volume and enhancing cooling and warming rates^16^.

Vitrification prevents ice crystal formation by rapidly cooling embryos to a glass-like state. This is achieved by briefly exposing embryos to high concentrations of cryoprotectant agents (CPAs), followed by immediate submersion into liquid nitrogen for ultrafast cooling^16,17^. In clinical practice, embryo vitrification is typically performed at the 4-cell, 8-cell or blastocyst stage (day 5-6 of development). These stages coincide with a critical period of genome-wide reprogramming in the embryo, including changes in methylation status, chromatin accessibility, degradation of maternal transcripts, and activation of the embryonic genome. While embryos appear morphologically normal after thawing, exposure to non-physiological conditions during this vulnerable period may result in cumulative negative effects.

Studies have shown that CPAs, while essential for vitrification, are also a direct source of toxicity. They have been linked to changes in chromatin architecture, DNA replication fork collapse, and DNA double-strand breaks, all of which contribute to genomic instability^18,19^.

Furthermore, embryo vitrification has been implicated in altering the expression of imprinting genes by disrupting DNA methylation levels and regulatory regions^20–25^. Impairment of these genes can directly influence fetal growth and offspring health^26,27^, underscoring the importance of the epigenetic landscape for normal mammalian development.

Cytoskeletal derangements have also been reported, including altered blastomere shape and size, spindle abnormalities and microtubules disassembly. These disruptions can interfere with major embryonic transitions and cell fate specification^28,29^.

In this study, we focused on the impact of vitrification on two cellular components whose interplay contributes to epigenetic remodeling and maintenance during embryo development: mitochondria and cytoplasmic lattices. Mitochondria serve as a primary interface between the environment and the epigenome, providing key metabolites necessary for epigenetic modifications during zygote genome activation (ZGA)^30,31^. Consequently, perturbations in mitochondrial structure and activity have the potential to significantly alter long-term embryo development by inducing heritable changes in the epigenome^31,32^. Notably, mitochondrial structure and function have been shown to be disrupted in post-thawed human endometrial tissue^33^ and reproductive cells of various species^34^ following cryopreservation, which further motivated our study. Regarding cytoplasmic lattices, these filamentous structures store essential proteins involved in DNA methylation and demethylation during embryo development, ensuring proper epigenetic reprogramming^35^. Although the impact of cryopreservation on CPLs remains unknown, studies have shown that disruption of lattices-associated proteins has been linked to severe impairments in ZGA, causing developmental delays and imprinting defects which are linked to imprinting disorders in humans^35,36^.

Using a mouse model and advanced imaging techniques, we investigated the effects of a widely used vitrification protocol on early embryo development, with a particular focus on mitochondrial structure and function as well as the integrity of cytoplasmic lattices. Our findings reveal that vitrification causes significant delays in early embryonic development, likely due to impaired mitochondrial activity and disruptions in cytoplasmic lattices abundance during the initial cell divisions.

## METHODS

### Animal Care and Ethical Approval

For this study, C57BL/6J(B6) female mice and CBA/J(CBA) male were housed in cages with free access to water and pelleted food in a temperature-controlled room, maintained under an artificial 12-h light/dark cycle.

All animal experiments conducted in this study were approved by the Stockholm Ethical Committee for Animal Research (19705-2022) and complied with the Directive 2010/63/EU of the European Parliament on the protection of animals used for scientific purposes. Animal care and experimental procedures were performed and monitored at Preclinical Laboratory 3 (PKL4) at Karolinska University Hospital, Huddinge, in accordance with accepted standards for humane animal care.

### Superovulation and mating

Superovulation was induced in 6-8-week-old female mice using a combination of pregnant mare serum gonadotropin (PMSG) and human chorionic gonadotropin (HCG). On Day 0, mice received an intraperitoneal injection of PMSG. Forty-eight hours later (Day 2), they were given HCG. Immediately after HCG injection, female mice were mated 1:1 with proven fertile male mice. Successful mating was confirmed the following day (Day 3) by checking for the presence of a vaginal plug. Female mice were sacrificed via cervical dislocation. See Supplementary Figure S3 for a schematic overview.

### Embryo collection and culture

From euthanized female mice, ovaries and associated oviducts were dissected. Zygotes were released from the ampullae into pre-warmed M2 medium and transferred to a hyaluronidase solution (HYASE™-10X, Vitrolife) to facilitate the removal of surrounding cumulus cells. Zygotes were then washed in M2 medium, cultured in pre-equilibrated EmbryoMax Advanced KSOM Medium (Sigma-Aldrich) covered with mineral oil (OVOIL™, Vitrolife), and incubated at 37°C with 5% O2 and 6% CO2. See Supplementary Figure S3 for a schematic overview.

### Embryo vitrification and warming

Embryos from each female were equally distributed in fresh and vitrified groups. In the vitrified group, 2-cell mouse embryos were cryopreserved using the Cryotop Method developed by Kitazato^12,16^.

Briefly, embryos were incubated in Equilibration Solution for 10-15 minutes. ES contains a lower concentration of permeable cryoprotectants, which helps the embryos lose water gradually while beginning to take up the cryoprotectants. After an initial shrinkage, embryos returned to their original size and were transferred into Vitrification Solution 1 for 30 seconds, followed by Vitrification Solution 2 for another 30 second. VS1 and VS2 contain higher concentrations of cryoprotectants and often include additional components, such as sucrose, which creates an osmotic gradient to promote further dehydration. These steps ensure complete dehydration and adequate cryoprotectant permeation before rapid cooling.. The embryos were then loaded onto the surface of the Cryotop strip with minimal volume and then plunged directly into liquid nitrogen.

For warming, the Cryotop strip was immersed in Thawing Solution 1 at 37°C for 1 minute until all embryos were released. Embryos were then transferred through a series of solutions at room temperature: Diluent Solution for 3 minutes, Warming Solution 1 for 5 minutes, and Warming Solution 2 for 1 minutes. These steps ensure controlled rehydration of the embryos and removal of cryoprotectants. Upon completion of the warming procedure, embryos were transferred to pre-equilibrated EmbryoMax Advanced KSOM Medium, overlaid with mineral oil, for further culture. Vitrified embryos were cultured for at least 2 hours to allow recovery before being subjected to additional experimental protocols. See Supplementary Figure S3 for a schematic overview.

### Time-lapse embryo culture

Fresh and vitrified 2-cell embryos were cultured individually in 16 microwells Geri dishes, allowing for the tracking of each embryo. They were then placed in a Geri incubator for 110-120 hours post-insemination. Bright field images were captured every 5 minutes at a resolution of 2pixels/µm, from the 2-cell stage until blastocyst stage. The Geri Connect and Assess 1.0 software was used to analyze all embryos, and the following morphokinetic parameters were annotated and compared between fresh and vitrified embryos: time intervals (in hours) from 2-cell stage to blastocyst stage.

To normalize the morphokinetic data, an initial normalization step was performed by defining the time of hCG injection as the starting point for both fresh and vitrified groups. An additional normalization step was applied to account for the delay caused by cryopreservation. Specifically, the time embryos remained frozen (approximately 2 hours) was subtracted from the total time each vitrified embryo took to reach each developmental stage. This ensured that only the time spent in culture after thawing was considered in the analysis.

### Immunofluorescence staining

Embryos were incubated at 37°C for 1 hour in 400nM Mitotracker Red CMXRos (Invitrogen; M7512) diluted in pre-equilibrated KSOM. After incubation, embryos were washed in KSOM for 1 hour at 37°C and then fixed in 4% paraformaldehyde (Sigma-Aldrich) for 20 min at RT. Embryos were then washed 3×5 minutes in PBA (PBS with 0.5% BSA, Sigma-Aldrich) at RT. Next, embryos were incubated in blocking solution (10% FBS in PBS, Sigma Aldrich) for 2 hours at RT or overnight at 4°C. Following blocking, embryos were incubated with Alexa Fluor™ 488 Phalloidin (Invitrogen; A12379) diluted 1:200 in blocking solution while protected from light, followed by 3×10 min washes in PBA. Embryos were mounted onto m-Slide 8-Well Glass Bottom (Ibidi) in droplets of VECTASHIELD Antifade Mounting Medium with DAPI (Vector Laboratories).

### Embryo imaging and analysis

Embryos were imaged using a Nikon Ti2 inverted spinning disk confocal microscope with 60× water objective, controlled by NIS-Elements software at the Live Cell Image Core Facility (LCI) of Karolinska Institutet. To ensure consistency and reproducibility, embryos were imaged using the same microscope settings, including laser intensity and exposure time. Z-stack (1μm steps) images were acquired sequentially for each channel along the embryo’s depth.

The acquired Z-stacks were opened in their native format in Imaris Software (Oxford Instruments version10.1), where 3D-reconstructions were automatically generated. Deconvolution of the MitoTracker channel was performed using a Gaussian filter to reduce noise. Subsequently, the images were converted to surface renderings, using the *Surfaces* model for Mitochondria and the *Cell* model for embryo blastomeres.

Imaris identified each individual mitochondrion as a surface by detecting MitoTracker fluorescence intensity through local contrast. The parameters for mitochondria surface rendering were manually adjusted for each embryo to ensure precise measurements, rather than relying on a fixed a threshold. Embryo blastomeres were rendered using the automatic Cell recognition function, based on the fluorescence intensity of the cell boundary stained with Phalloidin. When the automatic approach proved inadequate, manual rendering was rendered by profiling the Phalloidin signal every three Z-stack sections across the embryo’s length, using a vertex spacing of 2,76 µm.

Once surfaces were rendered, parameters such as mitochondrial volume and intensity, as well as blastomeres volume and area were generated for quantification. Mitochondrial volume refers to the proportion of an embryo blastomere volume occupied by mitochondria and was calculated as follows: total mitochondrial volume (µm^3^): divided by volume of the blastomere (µm^3^).

All images were prepared using the same brightness, contrast and color adjustments in Imaris.

### Transmission electron microscopy

Embryos were immersed in a fixative solution (2.5% glutaraldehyde buffered in 0.1M phosphate buffer pH 7.4) for 1 hour at RT and stored at 4°C. Fixed samples were then washed in 0.1M phosphate buffer, followed by post-fixation in 2% osmium tetroxide in 0.1M phosphate buffer, pH 7.4, at 4°C for 2 hours. Subsequently, embryos were dehydrated in ethanol followed by acetone and then resin infiltrated and embedded in LX-112 (Ladd Research). Ultrathin sections (~80-100 nm) were prepared using an Ultramicrotome EM UC7 (Leica) and transferred onto Formvar-stabilized slot grids. Sections were then contrasted with uranyl acetate followed by lead citrate. The slot grids were examined in a HT7700 transmission electron microscope (Hitachi High-Technologies,) at 80 kV, and digital images were acquired using a 2k x 2k Veleta CCD camera (Olympus Soft Imaging Solutions) at the Electron Microscopy Core Facility (EMiI) at Karolinska Institutet.

For cytoplasmic lattice quantification, a 4×4 µm region of interest (ROI) was selected within the cytoplasm, avoiding organelles that could obscure the lattices. This fixed ROI size ensured consistency across all analyzed images while being large enough to capture a sufficient density of cytoplasmic lattice fibers. Although the ROI dimensions remained constant, its placement was adjusted in each image to focus on cytoplasmic lattices while excluding irrelevant structures. Only images captured at 2 µm magnification were analyzed, as this magnification provided optimal clarity, making cytoplasmic lattice fibers easily distinguishable and countable. Additionally, multiple images from different regions of the embryo were analyzed at each developmental stage to account for potential spatial variations in cytoplasmic lattices distribution.

### QUANTIFICATION AND STATISTICAL ANALYSIS

The standard error of the mean was used for error bars for box charts. Statistical analysis was performed with GraphPad software (version 10.2.3). The unpaired Mann-Whitney U test was used to compare fresh and vitrified embryos. For all comparisons, P < 0.05 was considered to indicate statistical significance. On graphs, *P < 0.05, **P < 0.01, ***P < 0.001 and ****P < 0.0001. All data are from at least two independent experiments.

## RESULTS

### Time-lapse monitoring reveals notable delays in the developmental rhythm of vitrified mouse embryos

To compare the developmental kinetics between fresh and vitrified/warmed embryos, we used a time-lapse imaging system to monitor progression from the 2-cell stage to the blastocyst stage (Figure 1A). Vitrification and warming were performed at the 2-cell stage, a critical point in mouse preimplantation development that coincides with the onset of ZGA. This early stage also provides an opportunity to assess developmental progression and evaluate the impact of vitrification on early embryonic development within the limited timeframe of *in vitro* culture.

**Figure 1.**
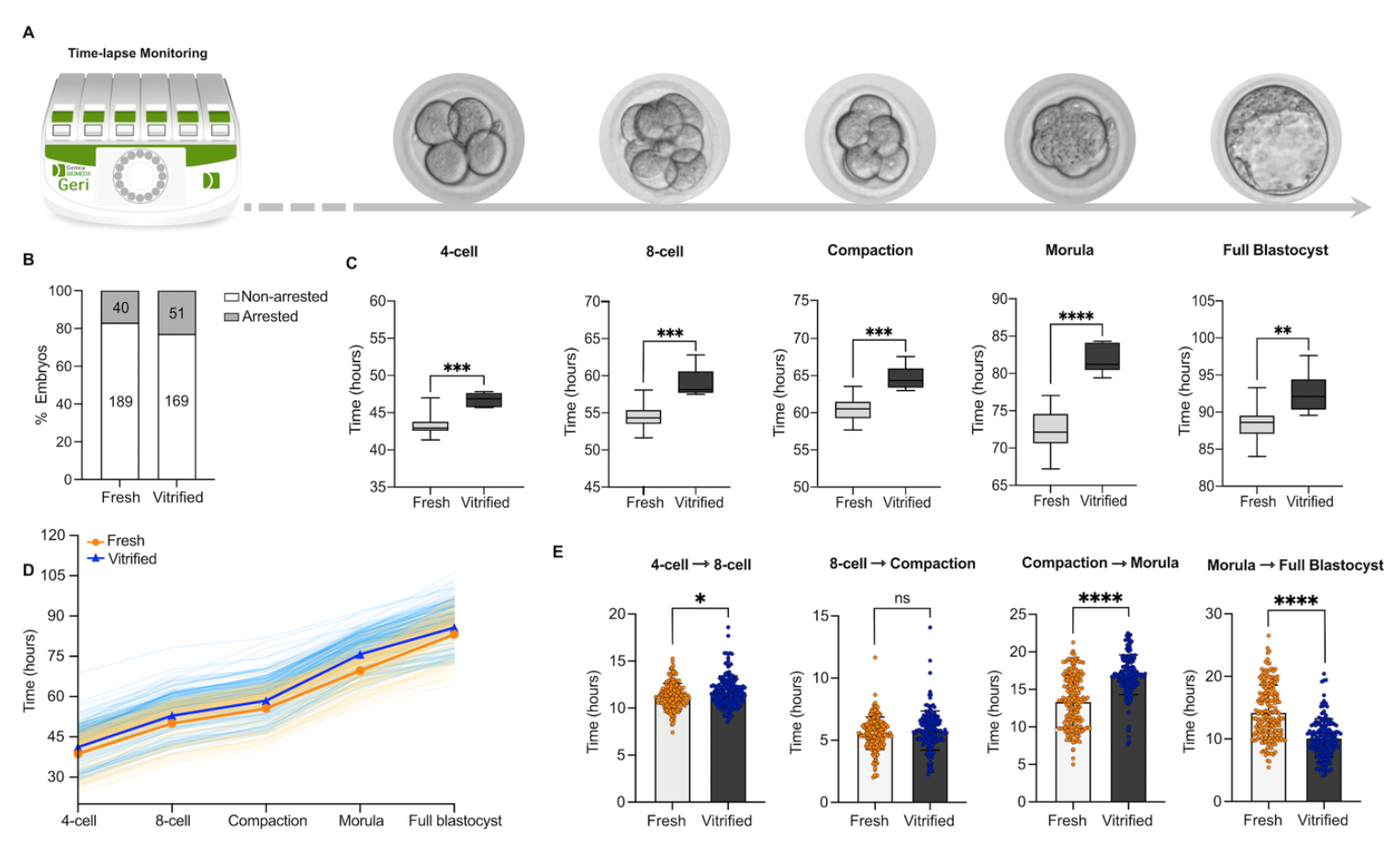
Time-lapse monitoring reveals notable delays in the development rhythm of vitrified mouse embryos. **(A)** Developmental progression of fresh and vitrified/warmed embryos was monitored using a GERI® time-lapse incubator. Embryos were vitrified and warmed at the 2-cell stage. Representative time-lapse images show the morphological appearance of mouse embryos at each developmental stage (left to right), captured at 10x magnification. **(B)** Percentage of non-arrested versus arrested embryos in fresh and vitrified/warmed groups. Embryonic arrest was defined as the absence of cellular division for at least 24 hours. **(C)** Time (in hours) for fresh and vitrified/warmed embryos to reach the 4-cell, 8-cell, compaction, morula, and full blastocyst stages. Results are presented as box-and-whisker plots, displaying median and interquartile ranges (boxes) and minimum and maximum values (whiskers). **(D)** An alternative visualization of the data presented in panel C, showing the developmental progression of individual embryos from the 4-cell to full blastocyst stage. Each line represents a single embryo: yellow lines indicate fresh embryos with the group average in orange; blue lines indicate vitrified/warmed embryos with the group average in dark blue. **(E)** Stage-specific durations (in hours) between transitions:4-cell to 8-cell, 8-cell to compaction, compaction to morula, and morula to full blastocyst. Each dot represents one embryo. Data are represented as mean ± SEM. Statistical comparisons in (C) and (E) were performed using unpaired two-tailed Mann-Whitney test: (ns) > 0.05, **p < 0.01, ^***^p < 0.001, ^****^p < 0.0001. The same set of embryos (229 fresh and 212 vitrified/warmed), derived from four independent experimental batches, was used for all analysis in panels A-E. Affinity Designer was used to create the illustration of the GERI® incubator.

We first evaluated the rate of embryonic arrest in fresh versus vitrified/warmed embryos. Embryos that failed to progress for at least 24 hours were considered arrested. Although a slight increase in embryonic arrest was observed in the vitrified/warmed group, this difference was not statistically significant when compared to the fresh group (Figure 1B).

Using a time-lapse Geri® incubator, fresh and vitrified/warmed embryos were individually tracked, and the exact time required to reach each developmental stage was recorded. Significant developmental delays were observed in vitrified/warmed embryos compared to fresh embryos. On average, delays ranged between 3-4 hours per developmental stage, with the most substantial cumulative delay (approximately 9 hours) observed by the time embryos reached the morula stage (Figure 1C). To further illustrate this developmental delay, the progression of each individual embryo is shown in an alternative format in Figure 1D.

To investigate whether the observed delay in vitrified/warmed embryos was a global developmental shift or stage-specific, we performed an interval-specific analysis (Figure 1E). For each embryo, we calculated the duration of key stage transitions: 4-cell to 8-cell, 8-cell to compaction, compaction to morula, and morula to full blastocyst. Vitrified/warmed embryos required, on average, one additional hour to transition from the 4-cell to 8-cell stage compared to fresh embryos (11.25 h vs. 11.58h). No significant difference was observed between groups for the 8-cell to compaction interval (5.58 h vs. 5.75 h). In contrast, vitrified/warmed embryos showed a substantial delay in the compaction to morula transition, taking nearly four hours longer than fresh embryos (13.42 h vs. 17.00 h). Interestingly, this trend reversed in the morula to blastocyst transition, where vitrified embryos developed more rapidly, reaching the blastocyst stage approximately three hours earlier than fresh counterparts (10.00 h vs. 13.50 h).

Despite these differences in developmental timing, blastocyst expansion was not visibly compromised by vitrification. As an indicator of developmental progression, the diameter of fresh and vitrified/warmed blastocysts was assessed, by measuring their maximum width (excluding zona pellucida)^37,38^. There was no significant difference in blastocyst diameter between groups, with average diameters of 104.7μm (fresh) and 103.9μm (vitrified/warmed) (Figure S1).

Taken together, although the overall developmental progression of vitrified embryos was delayed compared to fresh controls, interval-specific analysis revealed that the delay was not uniformly distributed across developmental stages. In particular, the accelerated transition from morula to blastocyst in vitrified/warmed embryo suggests that vitrification may alter the dynamics of specific transitions rather than simply shifting the entire developmental timeline.

### Vitrification alters morphometric parameters and might induce cell damage spots in 2-cell stage embryos

During the vitrification and warming process, embryos are exposed to fluctuations in osmotic pressure caused by the high concentrations of cryoprotectants used. In clinical settings, embryo quality and viability are typically assessed based on morphological characteristics including membrane integrity, blastomere survival and cytoplasmic appearance. In this study, we performed a quantitative analysis to compare fresh and vitrified/warmed embryos by evaluating morphometric parameters such as blastomere surface area and volume.

To achieve this, embryos were stained with Rhodamine-Phalloidin, an F-actin marker that reliably labels the cell membrane boundaries of blastomeres. Using Imaris Software, the surfaces of individual blastomeres were rendered, and 3D reconstructions are shown in Figure 2A.

**Figure 2.**
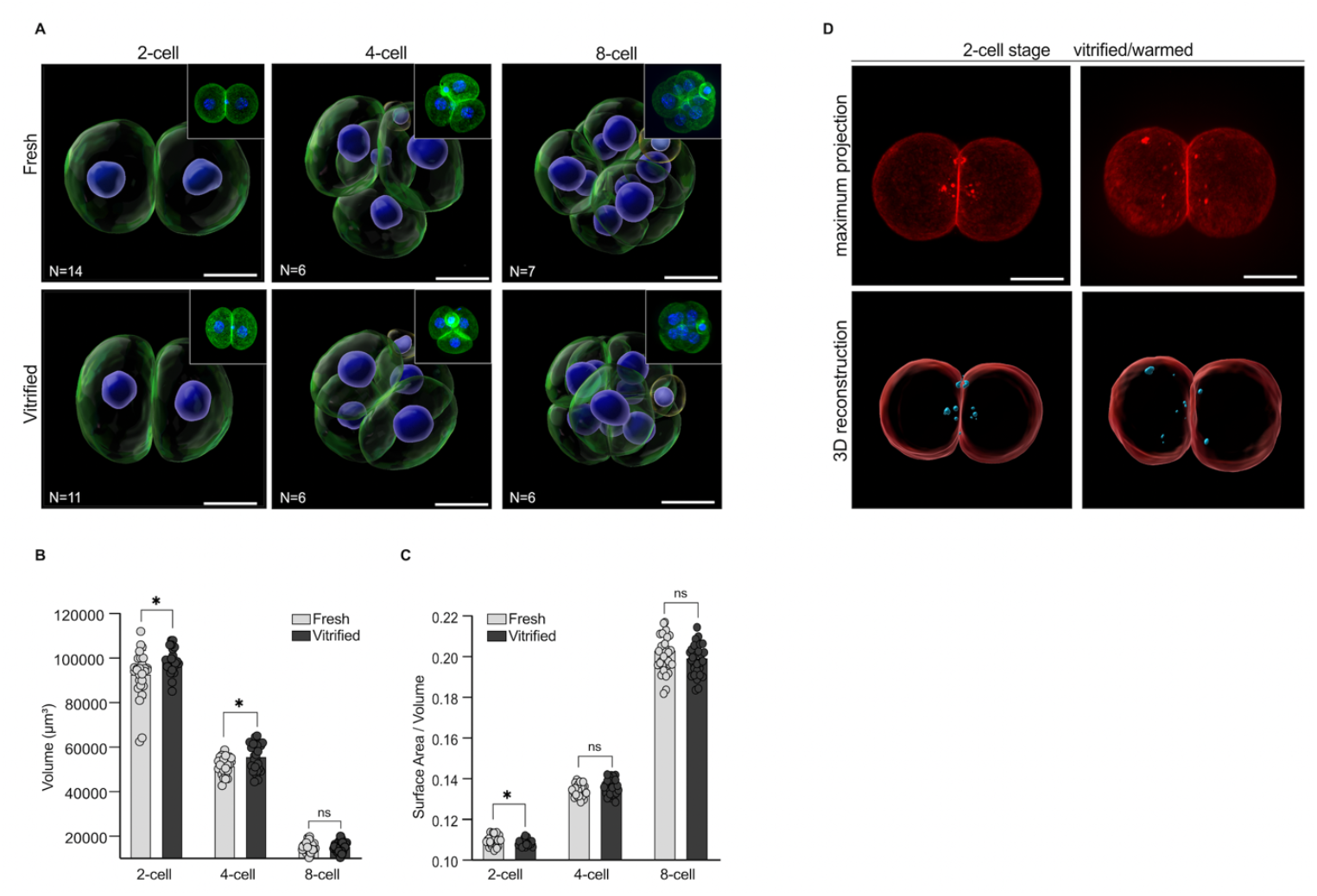
Vitrification alters morphometric parameters and may induce cell damage spots in 2-cell stage embryos. **(A)** 3D reconstructions of fresh and vitrified/warmed embryos labelled with Phalloidin (green) and DAPI (blue). Above each 3D reconstruction, a representative maximum projection of each embryo is shown (top right). **(B)** Graph representing blastomere volume (μm^3^). **(C)** Graph representing surface area-to-volume (S/V) ratio per blastomere. Each dot corresponds to one blastomere in a 2-cell, 4-cell or 8-cell embryo. Data are represented as mean ± SEM. Data were analyzed using unpaired two-tailed Mann-Whitney test: (ns) > 0.05, ^**^p < 0.01, ^***^p < 0.001, ^****^p < 0.0001. The number of embryos used is indicated in the figures. **(D)** Maximum projections and 3D reconstructions of fresh (not shown, N=10) and vitrified (N=8) 2-cell embryos labelled with Rhodamine-Phalloidin (red). F-Actin spots are highlighted in blue. Scale bars represent 10µm.

Like human embryos, mouse embryos exhibit an increase in the surface area-to-volume (S/V) ratio with each cell division, enhancing the efficiency of nutrient diffusion and waste removal. This developmental trend is observed in Figure 2B and 2C under fresh conditions. However, after thawing, 2-cell stage vitrified/warmed embryos exhibited a significant decrease in the S/V ratio compared to fresh embryos. This decrease likely reflects the effects of cryoprotectants, which are gradually diluted during warming to prevent osmotic shock. As the embryo rehydrates and begins to expand, its S/V ratio decreases.

By the 4-cell and 8-cell stages, the S/V ratios was comparable between fresh and vitrified/warmed embryos, highlighting the embryos’ capacity to adapt to environmental changes and restore homeostasis.

Interestingly, as shown in Figure 2D, we also observed multiple spots of increased fluorescent signal from Rhodamine-Phalloidin. These F-actin-rich spots were found exclusively in vitrified/warmed 2-cell embryos, located within the cytosol near the cellular membrane. Accumulation of actin has been reported as part of a stress response at sites of cellular damage, to trigger activation and recruitment of cellular repair mechanisms^39–41^. Based on these observations, we can only speculate that these actin-rich spots may represent a vitrification-induced stress response aimed at restoring the embryos’ cytoskeleton structure.

### Vitrification disrupts cytoplasmic lattices and mitochondrial structure during early mouse embryo development

For a detailed examination of the embryo’s cellular ultrastructure, transmission electron microscopy was applied on both fresh and vitrified/warmed embryos, at the 2-cell, 4-cell and 8-cell.

The first notable observation was the altered abundance and integrity of cytoplasmic lattices, in vitrified/warmed embryos. To quantify cytoplasmic lattices’ density, a region of interest (ROI) was defined, and the same ROI was used to analyze all electron microscopy images. As illustrated in Figure 3A, only cytoplasmic lattice fibers within the defined ROI were quantified.

**Figure 3.**
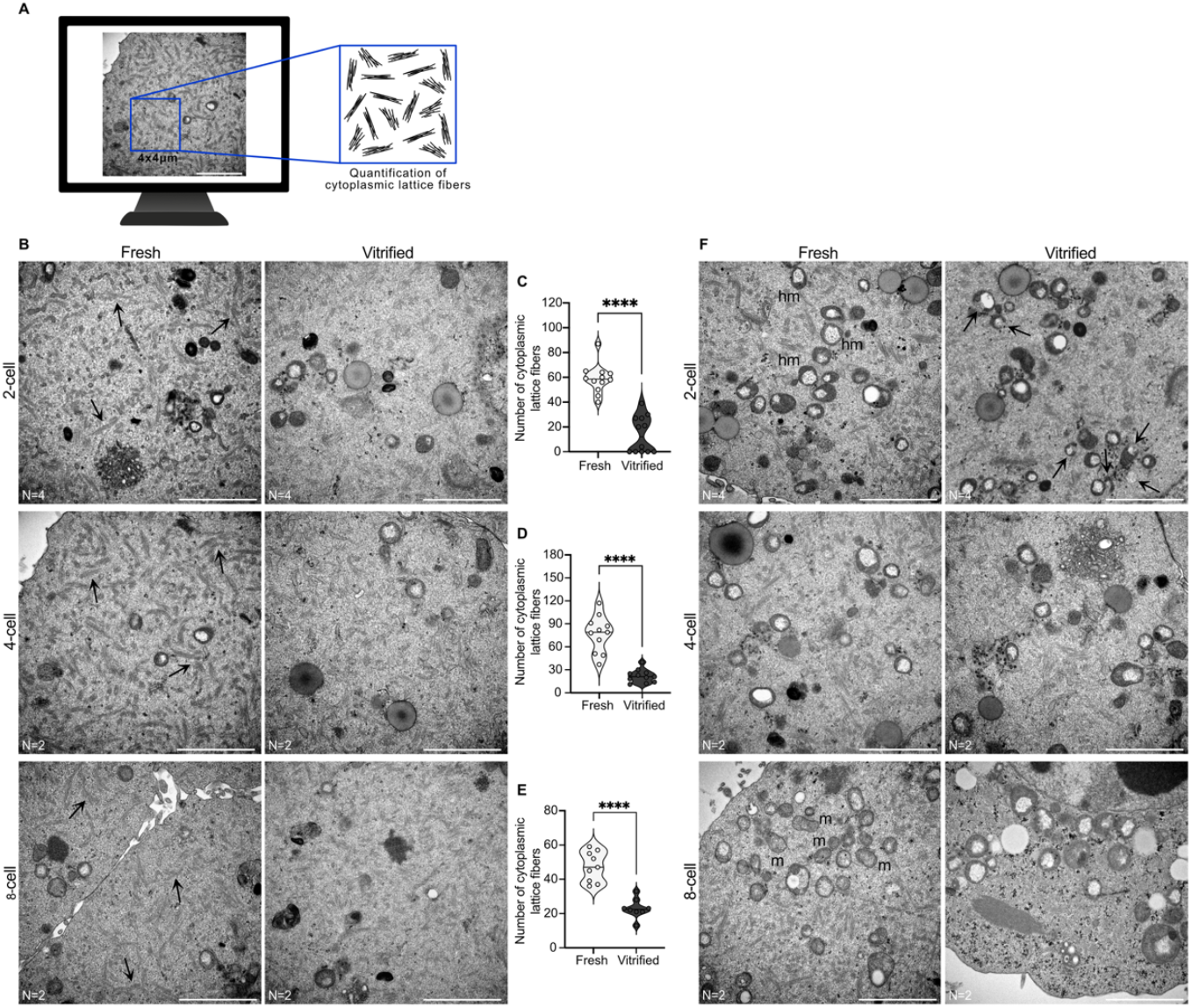
Vitrification disrupts cytoplasmic lattices and mitochondrial structure during early mouse embryo development. **(A)** Schematic representation of cytoplasmic lattices quantification within a 4×4µm region of interest (ROI). Depending on the developmental stage, micrographs from different regions of the embryo were analyzed: three micrographs for 2-cell stage embryos and six to eight micrographs for 4-cell and 8-cell stage embryos. **(B)** Representative transmission electron micrographs of fresh and vitrified/warmed 2-cell, 4-cell and 8-cell embryos, illustrating the distribution and abundance of cytoplasmic lattice fibers. **(C,D,E)** Quantification of cytoplasmic lattice fibers in fresh versus vitrified/warmed embryos at each developmental stage. Each dot represents one analyzed micrograph. Results are presented as violin plots, where the width represents the distribution of values, and the central line indicates the median. **(F)** Transmission electron micrographs of fresh and vitrified/warmed 2-cell, 4-cell and 8-cell embryos illustrating mitochondrial structure. Arrows indicate damaged mitochondria. Abbreviations: (hm) hooded mitochondria; (m) maturing mitochondria. Scale bars represent 2µm. Number of embryos analyzed are indicated below the micrographs. See also Figure S2.

At the 2-cell stage, the cytoplasm of fresh embryos was densely filled with cytoplasmic lattice fibers, while in vitrified/warmed embryos these fibers were significantly reduced or completely absent (Figure 3B and 3C). At the 4-cell stage, cytoplasmic lattices remained well-defined and visible in fresh embryos, while in vitrified/warmed embryos, they were significantly reduced and structurally altered, making individual filaments within the cytoplasmic lattices fibers difficult to distinguish (Figure 3B and 3D). By the 8-cell stage, cytoplasmic lattices remained evident in fresh embryos, but in reduced amounts compared to earlier developmental stages. In vitrified/warmed embryos, these lattices were further reduced and exhibited a less distinct structure (Figure 3B and 3E).

During early embryo development, hooded mitochondria are the predominant type^42,43^. Consistent with this, both fresh and vitrified/warmed embryos at the 2-cell stage displayed round-shaped mitochondria with a prominent vacuole-like space within the mitochondrial matrix, common characteristics of hooded mitochondria. Interestingly, in vitrified/warmed embryos, damaged mitochondria were observed with clear disruption of the mitochondrial outer membrane integrity (Figure 3F and Figure S2). At the 4-cell stage, hooded mitochondria remained the predominant type, and no significant structural differences were observed between fresh and vitrified/warmed embryos (Figure 3F). By the 8 cell-stage, fresh embryos displayed a higher number of matured mitochondria with a typical elongated structure. In contrast, most mitochondria from vitrified/warmed embryos appeared enlarged with highly distended vacuole spaces that occupied most of the mitochondrial volume, a potential sign of slow mitochondria maturation or degeneration (Figure 3F).

### Impact of vitrification on mitochondrial structure and cytoplasmic lattices in the blastocyst stage

Previous studies have reported poorer perinatal and obstetric outcomes following frozen blastocyst transfer^6,44^, suggesting that later-stage embryos may also be vulnerable to adverse effects from cryopreservation. Therefore, we also performed transmission electron microscopy on both fresh and vitrified/warmed blastocysts (day 5 of development). The vitrified/warmed group was further divided into two subgroups: blastocysts vitrified/warmed at the 2-cell stage and those vitrified/warmed at the blastocyst stage. This allowed us to investigate whether the timing of vitrification influences blastocyst quality.

Our first notable observation was the presence of electron-dense materials within the blastocele (Figure 4A). Initially, we suspected these could be artifacts from the preparation or sectioning process. However, since these materials were exclusively found within the blastocoel, we ruled out this possibility. While the exact nature of these electron-dense materials remains unclear, previous literature suggests they may be protein aggregates secreted by trophoblast cells to support blastocyst development^45–48^. Interestingly, these materials were observed in fresh blastocysts and those vitrified/warmed at 2-cell stage but were absent in blastocysts vitrified/warmed at the blastocyst stage.

**Figure 4.**
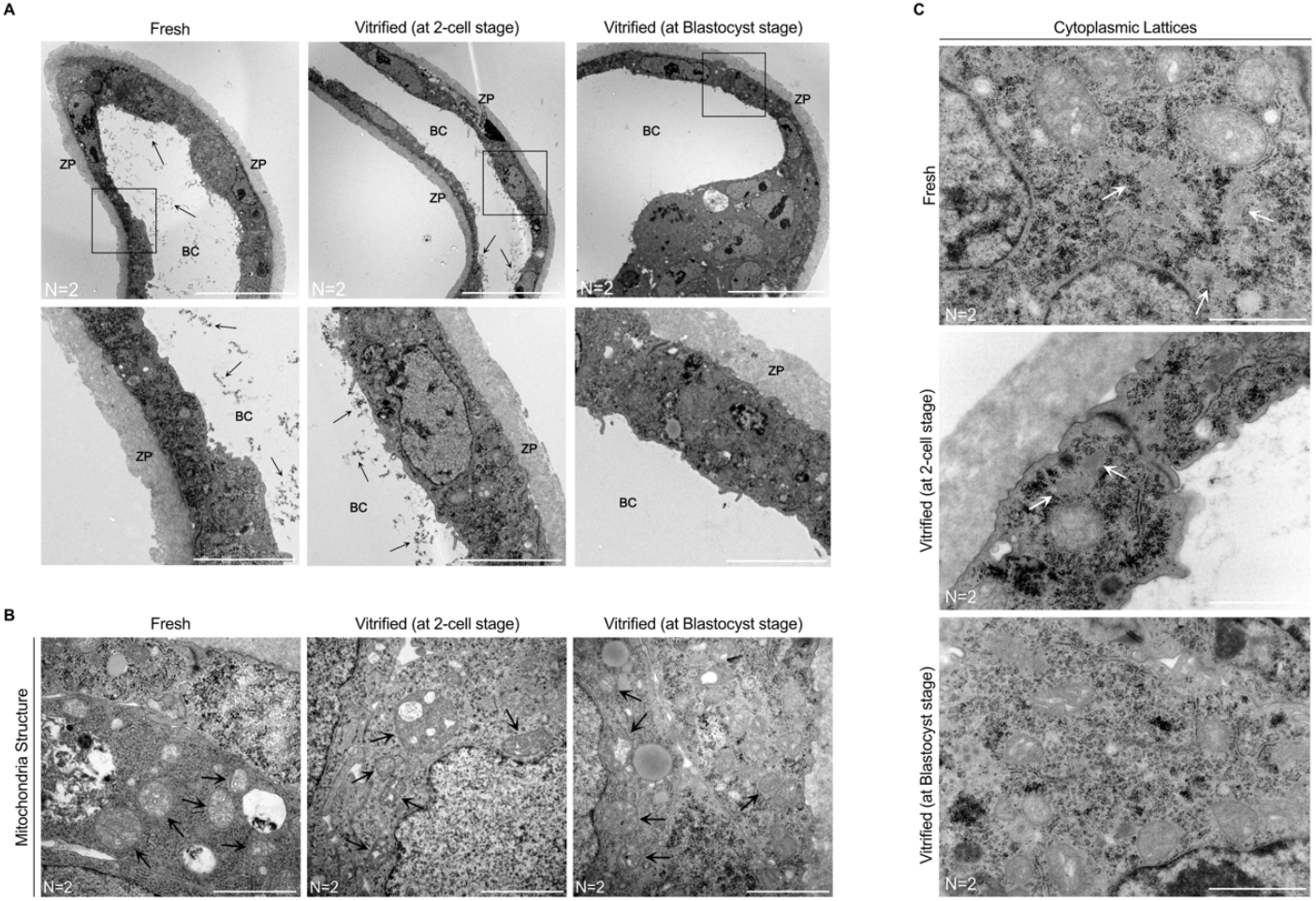
Impact of vitrification on mitochondrial structure and cytoplasmic lattices in the blastocyst stage. **(A-C)** Representative transmission electron micrographs of fresh blastocysts, blastocysts vitrified/warmed at the 2-cell stage, and blastocysts vitrified/warmed at the blastocyst stage. **(A)** Panels depict the presence (arrows) or absence of electron-dense materials within the blastocoel, with insets providing magnified views of the outlined regions. (**B)** Panels illustrate mitochondrial structure, with arrows highlighting well-developed cristae in fresh embryos and poorly developed cristae in vitrified/warmed embryos. (**C)** Panels show the presence (indicated by arrows) or absence of cytoplasmic lattice fibers. Scale bars represent **(A)** 20 µm (upper panel) and 5µm (lower panel), **(B, C)** 2µm. Number of embryos analyzed is indicated below each micrograph. Abbreviations: (ZP) zona pellucida, (BC) blastocoel.

Mitochondrial damage was observed in both vitrified/warmed subgroups (Figure 4B). Mitochondria in vitrified/warmed blastocyst exhibited poorly developed cristae and signs of vacuolization, indicative of compromised function and swelling. In contrast, fresh blastocysts displayed well-developed cristae, reflecting good mitochondrial integrity.

Cytoplasmic lattices were observed in fresh blastocysts, with well-organized and clearly defined filaments (Figure 4C). In blastocysts vitrified/warmed at the 2-cell stage, cytoplasmic lattices were also present, but in smaller agglomerates and at a lower abundance. In blastocysts vitrified/warmed at blastocyst stage, cytoplasmic lattices were not observable.

These findings highlight the impact of vitrification in mitochondria and cytoplasmic lattice integrity, even at a later stage of development. Moreover, our results suggest that the timing of vitrification, whether at the 2-cell stage or at the blastocyst stage, can have differential effects on blastocyst quality. Therefore, the timing of vitrification should be carefully considered to optimize the preservation of blastocyst integrity and minimize cryopreservation-induced damage.

### Vitrification disrupts mitochondria volume and membrane potential during early mouse embryo development

From ovulation until the morula stage in mammals, oocyte and early embryo development rely heavily on the mitochondria to meet their metabolic needs^49^. Variations in mitochondrial content and distribution have been associated with changes in embryo developmental rates in previous studies^50^. These variations are linked with adaptative mechanisms to help maintain a spatial balance of ATP supply in response to fluctuating energy demands during embryonic progression^51^.

Given the developmental delay observed in vitrified/warmed embryos compared to fresh, we aimed to assess how mitochondrial distribution and function are affected following vitrification/warming. To evaluate this, fresh and vitrified/warmed embryos at the 2-cell, 4-cell and 8-cell stage were stained with MitoTracker Red, a potential-dependent dye that accumulates in mitochondria based on their membrane potential.

In fresh 2-cell embryos, mitochondria were predominantly distributed in the perinuclear area and cytoplasm in a clustered pattern (Figure 5A and 5B). In contract, most vitrified/warmed embryos exhibited a more dispersed mitochondrial distribution within the cytoplasm (Figure 5A and 5B). The functional significance of mitochondrial clustering in early embryo development remains debated. Some studies suggest that clustering reflects an inability of embryos to develop *in vitro*^52^, while others link it to enhanced ATP production for rapid energy suply^42^. At later stages, 4-cell and 8-cell, both fresh and vitrified/warmed embryos displayed a similar mitochondrial distribution pattern, characterized by a dispersed pattern evenly distributed within the cytoplasm of each blastomere (Figure 5A).

**Figure 5.**
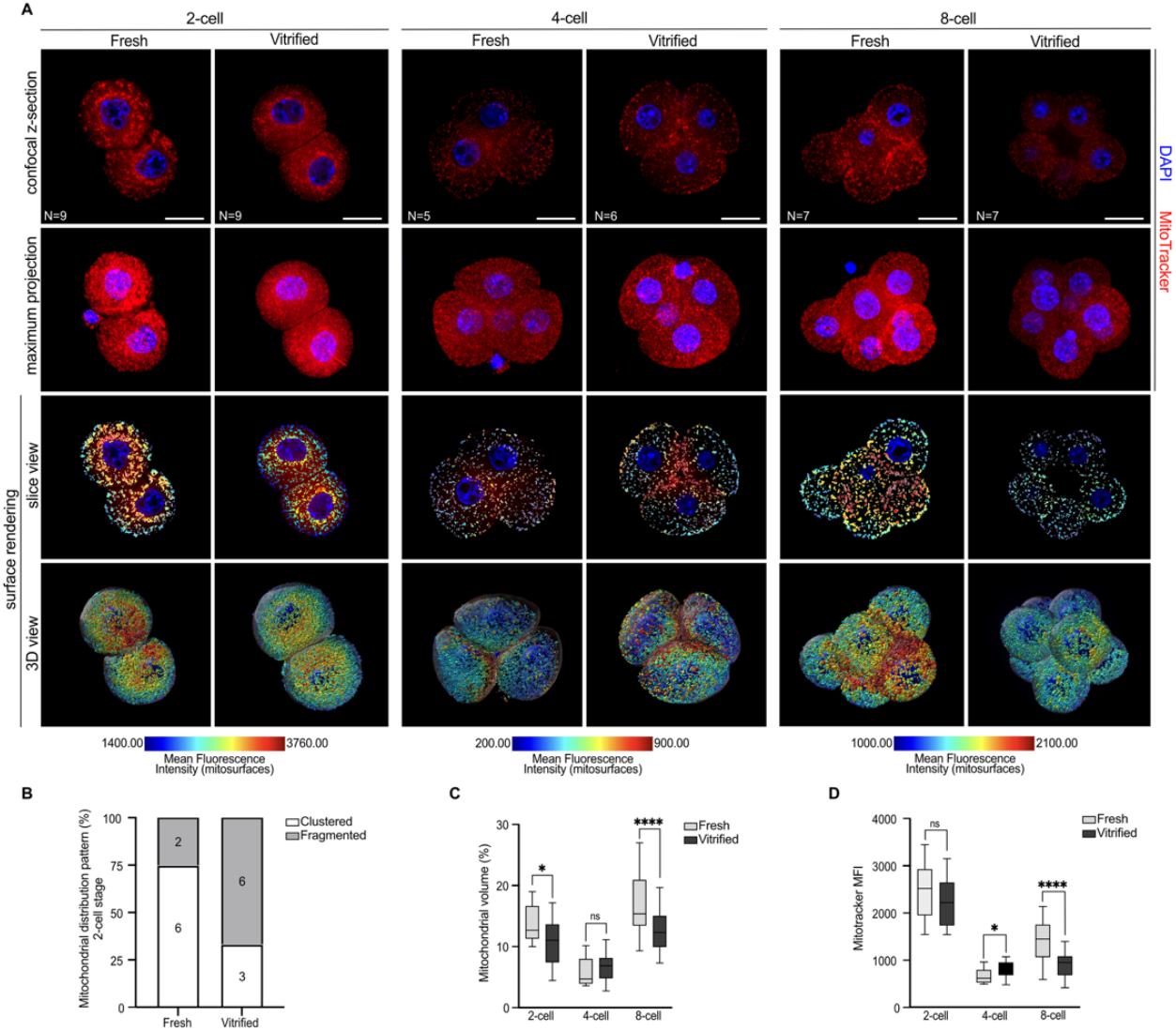
Vitrification disrupts mitochondrial distribution pattern, volume and membrane potential during early mouse embryo development. (**A**) Representative confocal z-sections (middle z-slice) and maximum projections (top two rows) of fresh and vitrified/warmed 2-cell, 4-cell, and 8-cell stage embryos, stained for mitochondria (Mitotracker, red) and DNA (DAPI, blue). Slice view mode and 3D reconstruction (third and fourth rows, respectively) illustrate rendered mitochondrial surfaces identified across a series of 60-90 optical sections. Mitotracker intensity is represented as a heat map, with a numerical color-coded scale below the panels. Scale bars represent 10µm. (**B**) Mitochondrial distribution pattern was evaluated in fresh and vitrified/warmed 2-cell stage embryos and classified as either clustered or fragmented. Numbers indicate the embryos classified under each category. Patterns in 4-cell and 8-cell embryos were identical for both fresh and vitrified/warmed conditions and thus were not included in the analysis. (**C,D**) Graphs showing mitochondrial volume (per blastomere) and Mitotracker fluorescence intensity (MFI, per blastomere) in fresh and vitrified/warmed embryos, respectively. Results are expressed as box-and-whisker plots, displaying median and interquartile ranges (boxes) and minimum and maximum values (whiskers). Data were analyzed using unpaired two-tailed Mann-Whitney test: (ns) > 0.05, ^**^p < 0.01, ^***^p < 0.001, ^****^p < 0.0001. The number of embryos analyzed is indicated in the first row of the panel.

To further investigate mitochondrial changes, Imaris software was used to generate surface renderings of mitochondria. This analysis identifies each individual mitochondrion or mitochondrial network as a separate rendered surface. The average mitochondrial volume per blastomere was calculated and is presented in Figure 5A, in both slice view and 3D view modes for fresh and vitrified/warmed embryos. At the 2-cell stage, vitrified/warmed embryos showed a significant decrease in average mitochondrial volume compared to fresh embryos. At the 4-cell stage, mitochondrial volume increased in vitrified/warmed embryos, though this change was not significant. Interestingly, by 8-cell stage, vitrified/warmed embryos showed a pronounced decrease in mitochondrial volume compared to fresh embryos (Figure 5C).

In addition to volume, mitochondrial fluorescence intensity was measured for each rendered mitochondrial surface, referred to as “mitosurfaces”. Heatmaps representing Mitotracker intensity are shown in Fig. 4A. At the 2-cell stage, Mitotracker intensity was comparable between fresh and vitrified/warmed embryos. However, at the 4-cell stage, Mitotracker intensity significantly increased in vitrified/warmed embryos compared to fresh, possibly suggesting a metabolic compensation by the embryos following vitrification. By 8-cell stage, Mitotracker intensity markedly decreased in vitrified/warmed embryos, consistent with the observed reduction in mitochondrial volume (Figure 5C and 5D).

Taken together, these findings demonstrate the potential consequences of vitrification on mitochondrial organization and membrane potential during early embryonic development.

## DISCUSSION

In this study, we contribute to the ongoing efforts in reproductive medicine to better understand the impact of embryo cryopreservation, with the goal of improving its safety for future pregnancies and newborns. Traditionally, cryopreservation techniques have been evaluated based on observable morphological changes and clinical outcomes, such as in vitro embryo survival, implantation, and birth rates. However, our findings highlights that such assessments may overlook subtle yet signification disruptions to early developmental processes.

To the best of our knowledge, this is the first study to provide a comprehensive comparison of developmental kinetics between vitrified/warmed and fresh mouse embryos, revealing significant differences across all pre-implantation stages. While vitrified/warmed embryos exhibited an overall developmental delay, stage-specific analyses showed that these delays were not uniformly distributed. This non-uniform pattern suggests that vitrification may not merely slow down overall development but instead, alter the timing and dynamics of specific morphogenetic events. One possible explanation for the accelerated morula-to-blastocyst transition in vitrified/warmed embryos is the activation of compensatory mechanisms. According to previous literature, mammalian embryos possess a degree of developmental plasticity that enables self-correction through gene expression adjustments, cytoskeleton reorganization, and flexible blastomere behaviour^53,54^. These adaptive processes may allow vitrified/warmed embryos to regain developmental synchrony by the morula stage. Alternatively, the acceleration may reflect vitrification-induced changes in cell adhesion and compaction dynamics. Proper morula formation relies on adhesion molecules, such as E-cadherin, and tight junction integrity^55–58^, both of which have been reported as disrupted in vitrified embryos of other species^59,60^. If compaction is incomplete or unstable, it may reduce the mechanical resistance needed for cavitation, leading to earlier blastocoel formation. In this scenario, the observed acceleration in blastocyst formation may not reflect true developmental progress but rather a shift in the timing of cavity formation due to altered cell behavior. Further investigation into adhesion molecules and junctional integrity is needed to confirm this possibility.

In this study, mouse embryos were vitrified using an open system commonly employed in fertility clinics, in which embryos embedded in cryoprotectant solution come into direct contact with liquid nitrogen.

Interestingly, distinct F-actin spots were observed near the cellular membrane exclusively in vitrified/warmed 2-cell embryos. F-actin accumulation has been previously reported to signal cellular damage, aiding repair at wound sites and in response to DNA damage^41,61^. A possible explanation for these F-actin spots, is a response to mechanical friction between the embryos and the vitrification strip device, causing localized damage and prompting an actin-based repair response. Alternatively, osmotic stress from the freezing and warming process could trigger F-actin reorganization. Osmotic stress and acute changes in cell volume are well-documented to induce cytoskeleton rearrangement and increase cellular F-actin in various cell types^62–66^. Notably, these F-actin spots were absent in later-stage vitrified embryos (4-cell and 8-cell stages), suggesting a cellular repair mechanism in response to vitrification at the 2-cell stage and, any potential damage was mitigated in subsequent developmental stages.

In this study, vitrification was performed at the 2-cell stage, coinciding with the major wave of ZGA in mice^67^. ZGA is a critical event in mammalian reproduction, during which the embryo assumes control of gene expression to regulate cell differentiation and further development. Failure to remodel the epigenome or activate embryonic transcription can result in implantation failure and developmental consequences, either *in utero* or later in life^67,68^. Successful reprogramming and embryo development rely on energy and metabolic cofactors produced by mitochondrial metabolism. As dynamic biosensors, mitochondria are highly sensitive to environmental perturbations, and their dysfunction have been shown to impair embryonic, fetal, and placental development^69,70^.

Early-stage embryos are enriched in “hooded” mitochondria, which are often described as “underdeveloped” or a “transition type mitochondria”, as they represent an intermediate stage in mitochondrial maturation^42,43,71–73^. These mitochondria provide initial glycolytic metabolic support, which transitions to oxidative phosphorylation as the embryo’s metabolic demands become more complex^42,71–73^. Indeed, hooded mitochondria were prevalent in 2-cell stage embryos, but vitrified/warmed embryos showed clear signs of mitochondrial membrane rupture. This likely reflects rapid temperature fluctuations and osmotic stress during vitrification/thawing, which can membrane lipid’s structure and composition, a common cause of cellular cryodamage^74^. Mitochondrial membrane rupture may allow leakage of critical mitochondrial components, impairing energy production. This aligns with the observed decrease in mitochondrial membrane potential at 8-cell stage (Figure 5A and 5D). Interestingly, vitrified/warmed 8-cell embryos also exhibited a higher number of hooded mitochondria compared to fresh embryos, potentially reflecting delayed mitochondrial maturation, or a broader developmental delay. Mitochondrial structure was also compromised at the blastocyst stage, regardless of whether vitrification occurred at the 2-cell or blastocyst stage. We observed mitochondrial swelling and poorly developed cristae, features similarly reported in vitrified blastocysts from sheep, porcine and bovine^75–77^.

Following fertilization, the early embryo relies almost exclusively on maternally inherited mitochondria, as de novo mitochondrial DNA replication does not occur until the blastocyst stage^78–81^. During cleavage divisions, mitochondrial content is partitioned into daughter cells without mtDNA replication^78–80,82,83^. Because mitochondrial biogenesis is minimal during these stages, any loss in mitochondrial content or quality cannot be readily compensated, supporting the existence of a functional “mitochondrial setpoint” essential for successful preimplantation development. Our findings indicate that vitrification at the 2-cell stage leads to an early mitochondrial insult, characterized by reduced mitochondrial volume, suggesting possible activation of mitophagy shortly after warming. By the 4-cell stage, we observed a significant rise in Mitotracker intensity in vitrified embryos. Although mitochondria biogenesis is limited during early development, embryos retain the ability to adjust the functional activity of their existing mitochondria. This includes metabolic reprogramming, upregulation of nuclear-encoded mitochondrial genes and alterations in mitochondrial dynamics^31,51,84,85^. The observed increase, at 4-cell stage, may represent a compensatory metabolic response wherein the embryo attempts to enhance mitochondrial function and energy output to support developmental progression despite earlier damage. However, the short duration of the 4-cell stage may limit the extent of this compensation. By the 8-cell stage both mitochondrial volume and Mitotracker intensity were markedly reduced in vitrified embryos, indicating a progressive decline in mitochondrial capacity. Further research is needed to clarify whether the observed changes in mitochondria reflect a metabolic adaptation or a cellular stress response to cryopreservation.

A remarkable finding of this study relates to cytoplasmic lattices. These highly ordered arrays of fibers, first identified in the 1960s^86,87^, have long remained enigmatic. Recent research by Shuch et al, demonstrated that mammalian oocytes store proteins essential for early embryonic development on cytoplasmic lattices^36^. Loss of these structures prevents the accumulation of these proteins and impairs early development, results in molar pregnancies or offspring with imprinting disorders^36^.

Our data revealed that cytoplasmic lattices in vitrified/warmed embryos were significantly reduced in abundance and exhibited compromised structural integrity during the initial cleavage stages. Since these structures store proteins essential for epigenetic reprogramming of the embryo, their impairment may offer a mechanistic explanation for epidemiolocal findings linking a higher incidence of imprinting disorders in children born from cryopreserved embryos^9^.

At the blastocyst stage, cytoplasmic lattices were still observed in fresh embryos, although in much lower abundance compared to earlier cleavage stages. While cytoplasmic lattices in blastocysts remain poorly studied, they are known to be dynamic structures that undergo extensive changes in spatial organization coinciding with major developmental transitions such as fertilization, blastomere compaction, and blastocyst formation^88–90^. Over time, as the embryo develops, these structures eventually fragment or disassemble^88–90^. This is consistent with the reduction in cytoplasmic lattices abundance observed in fresh blastocysts. This reduction likely occurs because most maternal RNA and proteins have been degraded or used, and the zygotic genome for transcription and protein synthesis now drives development.

In vitrified conditions, cytoplasmic lattices abundance was even lower or completely absent. Given that these structures were already observed in reduced quantities under fresh conditions, vitrification may have further degraded them. Whether this has a significant impact on blastocyst development and quality requires further investigation.

It is also important to mention that previous studies demonstrated that disruption of specific lattices-associated proteins can alter mitochondria morphology and distribution, as well as compromises energy production in mouse oocytes and zygotes^91–94^. However, the exact mechanisms by which cytoplasmic lattices contribute to mitochondrial function remains unclear. Further research is needed to determine whether disruption of cytoplasmic lattices plays a role in the observed mitochondrial dysfunction in vitrified embryos, and consequently, their developmental delays.

In conclusion, mitochondrial and cytoplasmic lattices alterations may underlie the disrupted developmental kinetics observed in mice vitrified/warmed embryos. Further studies are essential to validate these findings and to explore their translational relevance to human embryos. A deeper understanding of how cryopreservation affects subcellular structures, and developmental programming will be critical in refining assisted reproductive technologies and ensuring the long-term health of individuals conceived through these methods.

### Limitations of the study

It is worth noting that, although our work provides intriguing insights into the impact of cryopreservation on embryo development, several issues remain to be addressed.

Firstly, our research was conducted using a mouse experimental model, which may not fully translate to humans due to considerable species differences in embryonic development, metabolism, and timings of key events such as ZGA. Additional research is needed to validate our findings and determine their relevance to human embryos.

Part of our mitochondrial assessment was performed on fixed embryos using confocal microscopy. While informative, this approach limits the ability to assess mitochondrial real-time behavior, as these organelles are highly responsive to the cell cycle, metabolic state and environmental conditions. Although embryos were tightly time-controlled in this study, live imaging approaches will be essential in future research to more accurately capture mitochondrial dynamics during development. In addition, although Mitotracker is widely used to evaluate mitochondrial membrane potential, it provides only a partial picture of mitochondrial function. Future studies should incorporate complementary approaches using additional dyes, such as JC-1 for more precise membrane potential measurement or functional assays for ATP quantification for a more comprehensive view of mitochondrial health and activity in cryopreserved embryos.

Lastly, further structural studies are needed to explore the three-dimensional organization of cytoplasmic lattices after cryopreservation. Although, transmission electron microscopy provided valuable details, its two-dimensional nature may result in the loss of important information along the embryo’s depth. In addition, due to technical demands and time-intensive nature of transmission electron microscopy, our sample size for ultrastructural analysis was limited.

## RESOURCE AVAILABILITY

### Lead contact

Requests for further information and resources should be directed to and will be fulfilled by the lead contact, Arturo Reyes-Palomares.

### Materials availability

This study did not generate new unique reagents.

### Data and code availability

- This paper does not report original code. All data associated with this study are present in the paper or supplemental information.
- Any additional information required do reanalyze the data reported in this paper is available from the lead contact upon request.

## ACKNOWLEDGMENTS

We would like to express our sincere gratitude to the staff at the Animal Facility, Live-Cell Imaging Facility, Biomedicum Imaging Core Facility, and Electron Microcopy Core Facility of Karolinska Institutet for their invaluable technical assistance. In particular, we extended our thanks to Muntaha Fartoo for her assistance with animal experiments and embryo collection; Gabriela Imreh for her support with the embryo immunofluorescence protocol and confocal microscopy; Göran Månsson for his expertise and advice on confocal image analysis, particularly with Imaris software; and Dr. Lars Haag and Lisa Sjöwall for their help with embryo processing for transmission electron microcopy and image collection. We also wish to acknowledge Birgitta Lindqvist for her support in managing laboratory resources, handling orders, and ensuring proper management of the laboratory environment. The research leading to these results was funded by KI Research Foundation Grants 2024-2025(2024-02566), Vetenskapsrådet Research Grants Open call 2023 (Medicine and Health) 2023-01872. The current work of ARP is supported by the Spanish Ministry (Beatriz Galindo Program).

## AUTHOR CONTRIBUTIONS

Conceptualization, M.T.B., K.A.R.W., and A.R.P.; methodology, M.T.B. and A.R.P.; investigation, M.T.B., K.D., J.A.R.M., V.S.; formal analysis, M.T.B., K.D., J.A.R.M., V.S, and J.Z.; writing—original draft, M.T.B.; writing - review and editing, M.T.B., K.A.R.W., and A.R.P.; visualization, M.T.B; funding acquisition, K.A.R.W. and A.R.P.; supervision, K.A.R.W. and A.R.P. All authors read and revised the manuscript.

