## Supplementary Material for "Vitrification-warming delays preimplantation development and impairs mitochondrial function and cytoplasmic lattices integrity in mouse embryos"

### Supplemental Figures

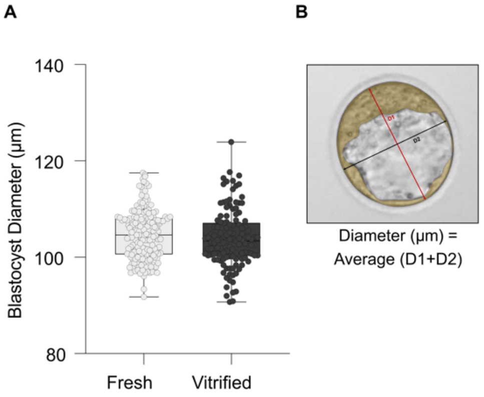


**Figure S1. Measurement of blastocyst diameter (µm) between fresh and vitrified/warmed groups, related to Figure 1. (A)** Graphical representation of the diameters of fresh (N=229) and vitrified/warmed (N=212) blastocysts. **(B)** Schematic representation of the diameter measurement. The blastocyst diameter was calculated by averaging two cross-sectional measurements (D1 + D2), excluding zona pellucida. Measurements were performed on time-lapse microscopy images at the time each embryo reached fully expanded blastocyst stage.

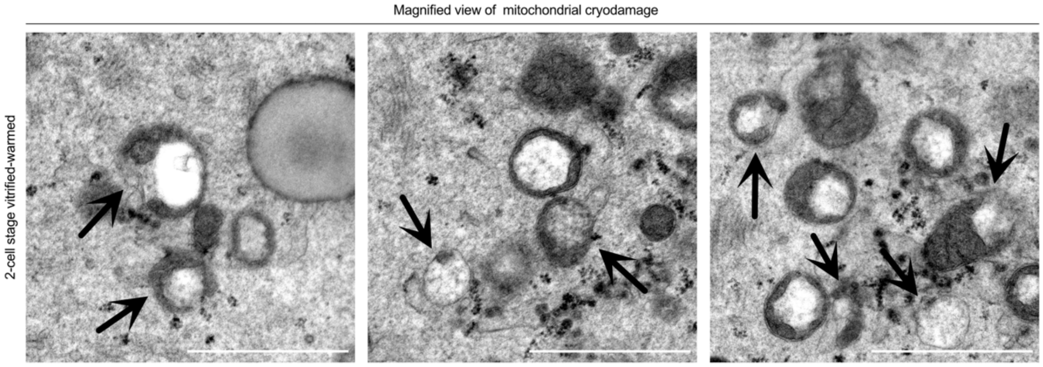


**Figure S2. Mitochondrial membrane damage in vitrified-warmed 2-cell stage embryos, related to Figure 3.** Representative magnified transmission electron micrographs showing mitochondrial ultrastructure in 2-cell stage embryos following vitrification and warming. All micrographs are cropped regions of interest obtained from a larger stitched TEM image acquired at 15,000x magnification. Black arrows indicate damaged mitochondria, characterized by disrupted membranes. Scale bars represent 1µm.

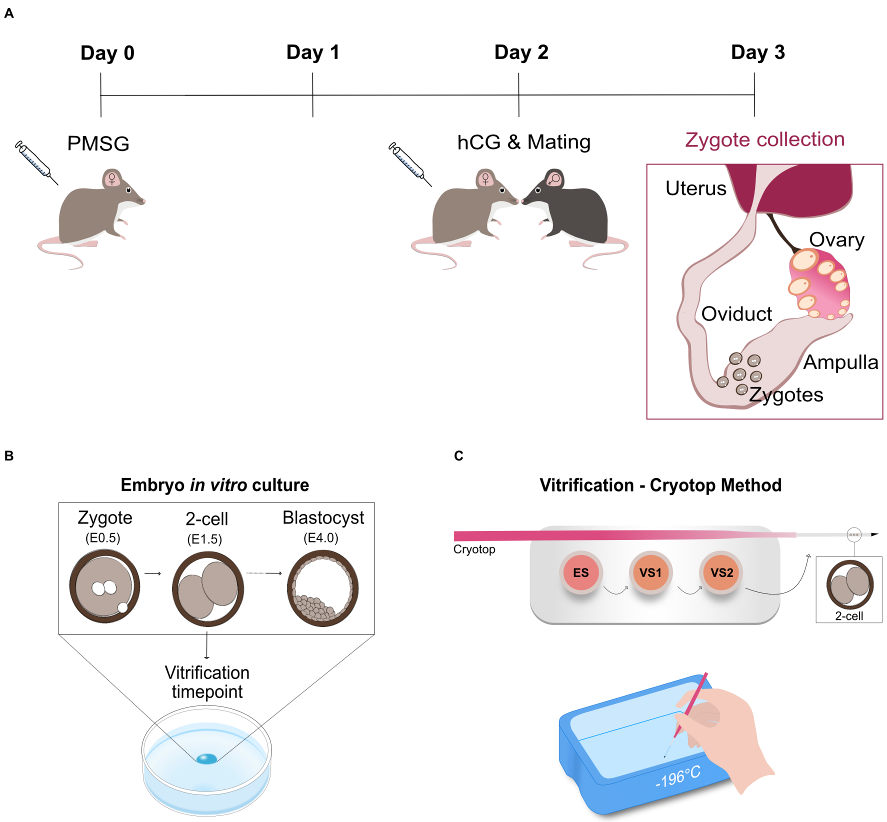


**Figure S3. Experimental setup, related to STAR Methods. (A)** Schematic timeline of mice superovulation, mating and zygote collection. Superovulation was induced with pregnant mare serum gonadotropin (PMSG) on Day 0, followed by human chorionic gonadotropin (hCG) on Day 2. Following hCG injection, females were mated with males. Zygotes were collected from the ampulla on Day 3 after dissection of the ovaries and oviducts. **(B)** Zygotes (E0.5) were cultured in vitro to the blastocyst stage (E4.0). Embryos were vitrified and warmed at the 2-cell stage (E1.5), as indicated. **(C)** Schematic representation of the vitrification procedure using the Cryotop method. Embryos were first incubated in Equilibration Solution (ES) for gradual dehydration and initial cryoprotectant exposure. Embryos were sequentially exposed to Equilibration Solution (ES), then to Vitrification Solution 1 (VS1) and Vitrification Solution 2 (VS2), which contain increasing concentrations of cryoprotectants. Embryos were then loaded onto a Cryotop strip and immediately submerged in liquid nitrogen (–196 °C). Affinity Designer was used to create the illustration.
